# Imaging forward and Reverse Traveling Waves in the Cochlea

**DOI:** 10.1101/348847

**Authors:** A. Zosuls, L. C. Rupprecht, D. C. Mountain

## Abstract

The presence of forward and reverse traveling wave modes on the basilar membrane has important implications to how the cochlea functions as a filter, transducer, and amplifier of sound. The presence and parameters of traveling waves are of particular importance to interpreting otoacoustic emissions (OAE). OAE are vibrations that propagate out of the cochlea and are measureable as sounds emitted from the tympanic membrane. The interpretation of OAE is a powerful research and clinical diagnostic tool, but OAE use has not reached full potential because the mechanisms of their generation and propagation are not fully understood. Of particular interest and deliberation is whether the emissions propagate as a fluid compression wave or a structural traveling wave. In this study a mechanical probe was used to simulate an OAE generation site and optical imaging was used to measure displacement of the inner hair cell stereocilia of the gerbil cochlea. Inner hair cell stereocilia displacement measurements were made in the radial dimension as a function of their longitudinal location along the length of the basilar membrane in response to a transverse stimulation from the probe. The analysis of the spatial frequency response of the inner hair cell stereocilia at frequencies near the characteristic frequency (CF) of the measurement location suggests that a traveling wave propagates in the cochlear partition simultaneously basal and apical (forward and reverse) from the probe location. The traveling wave velocity was estimated to be 5.9m/s - 8m/s in the base (near CF of 29kHz - 40kHz) and 1.9m/s - 2.4m/s in the second turn (near CF of 2kHz - 3kHz). These results suggest that the cochlear partition is capable of supporting both forward and reverse traveling wave modes generated by a source driving the basilar membrane. This suggests that traveling waves in the cochlear partition contribute to OAE propagation.

## Introduction

The mammalian cochlea has remarkable frequency selectivity and dynamic range in part due to the physically distributed filtering (1) and a massively parallel nonlinear transduction system (2). To understand its function as a physically distributed (spatial) filter, an intimate knowledge of how information propagates along the length of the cochlea is paramount. For example, the use of otoacoustic emissions (OAE) as an investigative and diagnostic tool relies on understanding the modes of propagation of information both forwards and backwards in the cochlea.

OAEs are sounds emitted from the ear that originate as hydro mechanical perturbations in the cochlea (3). They can be measured noninvasively thus are an effective clinical diagnostic tool whose utility will be greatly increased once the origins and propagation mechanisms are fully understood. Theoretical models and experiments have identified wave reflection and nonlinear processes inside the cochlea as the primary sources of OAE generation(4).

The notion of traveling waves in the mammalian cochlea has been a debated topic in cochlear mechanics since the 1940s when they were measured by von Bekesy (5). The presence of a forward traveling wave mode in the cochlear partition (CP) is expected from the large body of work (5–9) where it was predicted and measured. However, there are contradicting results and much less evidence regarding the presence of reverse traveling waves in the cochlear partition. Data and analysis by He (10) suggest there is a forward propagating traveling wave along the CP but no reverse traveling wave (11). He’s experiments in gerbil (10) support the theory that the reverse wave propagates as a fast fluid compression wave in the scalae. Model results (12–15) and experimental results (16–18) support the theory of a slow reverse traveling wave as a propagation mechanism for OAEs. A slow wave refers to one in which the cochlear fluids and the CP impedance interact and support a mode of propagation that is transverse to the CP (figure1). A fast wave is the compression wave that results from sound propagating through the fluid on the scalae.

**Figure 1:**
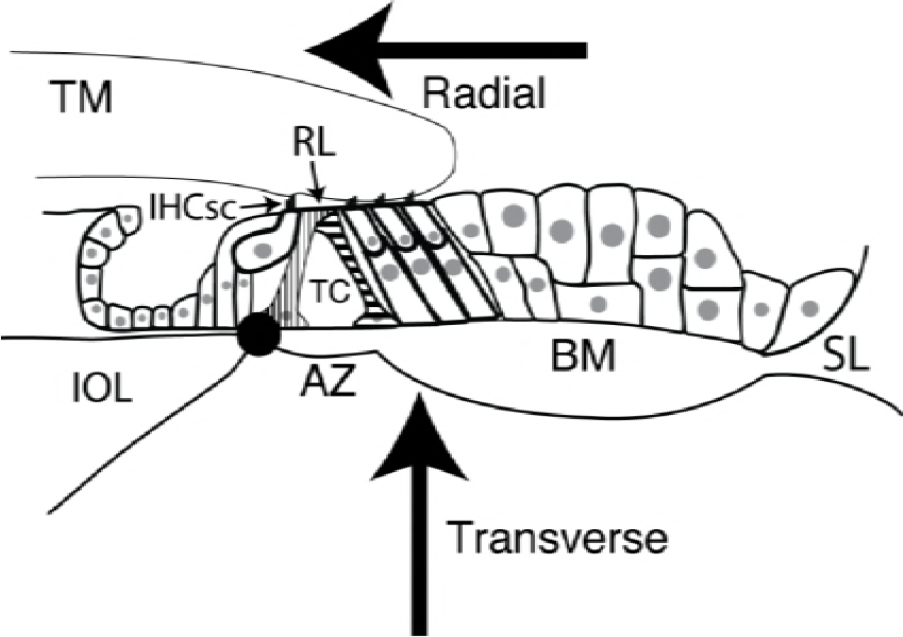
Illustration of a cross section of the cochlear partition (CP) in gerbil. The basilar membrane (BM) spans from the inner osseous lamina (IOL) to the spiral ligament (SL). The tunnel of Corti (TC) comprises the triangle formed by the inner pillar cell (vertical stripes), outer pillar cell (horizontal stripes), and a section of the BM called the arcuate zone (AZ). The AZ connects the bottom feet of the two pillar cells. The reticular lamina (RL) is the surface where the hair cell heads and pillar cells connect. In this study we measure the displacement of the inner hair cell stereocilia (IHCsc). The black dot represents the theoretical axis of rotation of the tunnel about the inner pillar cell foot. In this paper transverse displacement is in the direction normal to the plane of the BM. Positive transverse displacement of the CP is toward the scala vestibuli as show by the transverse arrow. Radial is defined as the dimension that spans the width of the BM, from the IOL to the SL. As seen in the figure, positive radial displacement is in the direction of the radial arrow (modiolar). Longitudinal is defined as the dimension spanning from base to apex and is normal to the plane of the figure on the page.

Basilar membrane (BM) velocity and displacement measurements are often used to investigate the mechanics and propagation modes of the cochlea, and how neural output relates to input from the middle ear. The BM consists of collagen fibers and is the mechanical foundation of the CP; the sensory and support cells are attached to it. Many measurements of BM displacement or velocity were recorded in the transverse direction (figure1) (2, 5, 19, 20). While providing biomechanically correct information regarding how the BM moves, these measurements should be regarded in context with the usual function of the cochlea’s biological sensors that innervate the auditory nerve: The inner hair cells (IHC) sense radial, not transverse displacement. The IHC membrane potential changes as a function of the displacement of their stereocilia (21). The location and orientation of the IHC stereocilia (IHCsc) suggest that a shearing motion of the IHC bodies relative to the tectorial membrane is the preferential method of stimulation (22). The shearing motion is primarily in the width (radial) dimension, although it has been shown that the CP vibrates in the height (transverse) dimension when stimulated with an acoustic pressure source from the stapes footplate (20, 23). It has been proposed that the transformation from transverse to radial displacement is a result of the pillar cell triangle rotating about the foot of the inner pillar cell in response to pressure across the basilar membrane (22, 24, 25) (figure 1).

The transverse direction was often measured because of theoretical and experimental evidence that to a rough approximation, the complex anatomy of the CP can be simplified to a tuned mechanical plate or beam, and the tuning parameters (impedance) can be deduced by knowledge of the pressure across and velocity of the CP (26).

In this study we investigated the radial displacement of the IHCsc as a function of their longitudinal location along the BM in response to transverse point displacement applied to the BM. Measuring radial displacement of the IHCsc as opposed to transverse displacement yields results in the dimension where the IHCsc are most sensitive. The ability to simultaneously measure the radial displacement of many IHCsc over a longitudinal section of the cochlea reveals wave propagation modes in this dimension. By using a point driving source that is local to the area being measured it is possible to investigate how waves propagate in the cochlear partition.

## Methods

Our imaging system is an evolution of the methods used by Karavitaki (27) to track outer hair cells. It enabled simultaneous measurement of displacement in the plane of the basilar membrane (longitudinal-radial plane) over approximately a 300 μm by 200 μm field of view. This facilitated simultaneous measurement of many IHCs and other structures, allowing a spatial displacement response to be extracted. The system consisted of an inverted microscope with a custom stroboscopic imaging system linked to a computer controlled probe used to mechanically stimulate the basilar membrane in the transverse direction. Experiments were conducted with sinusoidal stimulus frequencies ranging from 10 Hz to 42.5 kHz. This frequency range included the most sensitive frequencies (CF) for both measured longitudinal locations. Using digital image processing, subpixel displacements down to 8nm were resolved.

### Tissue preparation

Experiments were performed according to IACUC-approved procedures at Boston University on cochleae extracted from 29 - 59g female gerbils (*Meriones unguiculatus*). The animals were anesthetized with an intraperitoneal injection of Ketamine (80mg/kg dose) and Xylazine (20mg/kg dose). Once there was a full cessation of toe withdrawal and corneal reflexes, the animals were decapitated and the bullae removed and immersed in oxygenated artificial perilymph at room temperature. The perilymph (27, 28) consisted of a chloride modified solution composed of 140 mM D-GlcA, 6.6 mM NaCl, 100μM CaCl_2_, 2 mM KCl, 5 mM NaH_2_PO_4_, 100 μM MgCl_2_, 5 mM D-Clc, 5mM HEPES (298 mOsm, pH 7.3 adjusted with 1M NaOH). The cochlea was dissected out of the bulla and the stapedial artery was drained and removed to prevent red blood cell proliferation into the scalae.

In turn 1 preparations, the first turn of the cochlea (the base) was separated from the other two turns by carefully perforating the bone encompassing the scala vestibuli (SV) with a scalpel and then cleaving the scalae to expose the cochlear partition. This technique maintained the integrity of the stria and Reissener’s membrane. The round window was removed with a hook formed from the fine tip of a bent tungsten probe. Turn 1 preparations were stimulated through the round window aperture or through an enlargement of the round window boney aperture. The enlargement was made on the apical side of the round window by perforating the scala tympani (ST) and breaking bone fragments free with forceps. In turn 1 preparations, the distance from the base to the probe tip was measured optically during each experiment using a low magnification image of the preparation. The CF estimation was calculated using the frequency - distance place map from Muller (29).

The turn 2 preparations were separated from both turns 1 and 3. The first turn was separated from the second turn by perforating the first turn SV with a scalpel and then using the scalpel to cleave apart turns 1 and 2. This left the boney partition between the turn 1 SV and turn 2 ST intact. Next a portion of turn 3 was removed using a scalpel and forceps to expose the partition between the turn 3 ST and turn 2 SV. Once turn 2 was isolated, windows were made for the probe and optical path in both scalae by perforating and lifting out the boney partitions to expose the CP. The Muller place map (29) was used to estimate the CF by making a composite measurement of the lengths of the BM in 3 cochlea. The BM length was measured optically for each piece of the cochlea that was basal to the probe location. The segments were summed to make a composite length estimate that resulted in a CF estimate.

During dissection and preparation the cochlea was continuously immersed in room temperature oxygenated artificial perilymph. Frequent dissecting dish and culture medium exchange helped to maintain the health of the preparation and reduce the contamination of the preparation with bone and cellular debris. After the dissection was complete the preparation was bonded to a fixture with Pro CA cyanoacrylate adhesive and accelerator (Great Planes, Champaign, Illinois). A micropipette was used to dispense the adhesive and accelerator to minimize the possibility of seepage into the scalae during bonding. The fixture consisted of a stainless steel rod connected to a custom-made multi degrees of freedom fixture that was magnetically anchored to the microscope stage. An FD35 culture dish (World Precision Instruments, Sarasota, Florida) with a glass bottom for optical coupling to the water immersion objective was used to hold oxygenated culture medium which was changed throughout the experiment to maintain the quality of the tissue.

### Optics and stimulation

The cochlea-bar fixture was mounted on the stage of an IX 70 inverted microscope (Olympus Waltham, Massachusetts) with the preparation fully immersed in oxygenated culture medium with the scala vestibuli facing the objective (down) (figure 2A). The fixture was adjusted to align the cells being measured within the cochlear partition to the focal plane of the objective. All turn 1 experiments ranged from 1000 to 1600 μm from the base not including the length of the hook region. Turn 2 experiments were performed one turn apical to the base or approximately 7000 μm from the base. The stimulating probe was oriented normal to the basilar membrane. It was positioned longitudinally in the center of the field of view and radially under the outer pillar cell foot. The probe was brought into contact with the basilar membrane using a DC3001 motorized micromanipulator (World Precision Instruments, Sarasota, Florida) by advancing the probe in 2 μm increments until a slight change in focus of the pectinate region radial collagen fibers indicated they were displaced. Care was taken to prevent the probe from visually damaging or buckling the preparation. Once contact was established, the microscope was focused on the inner hair cell stereocilia.

**Figure 2.**
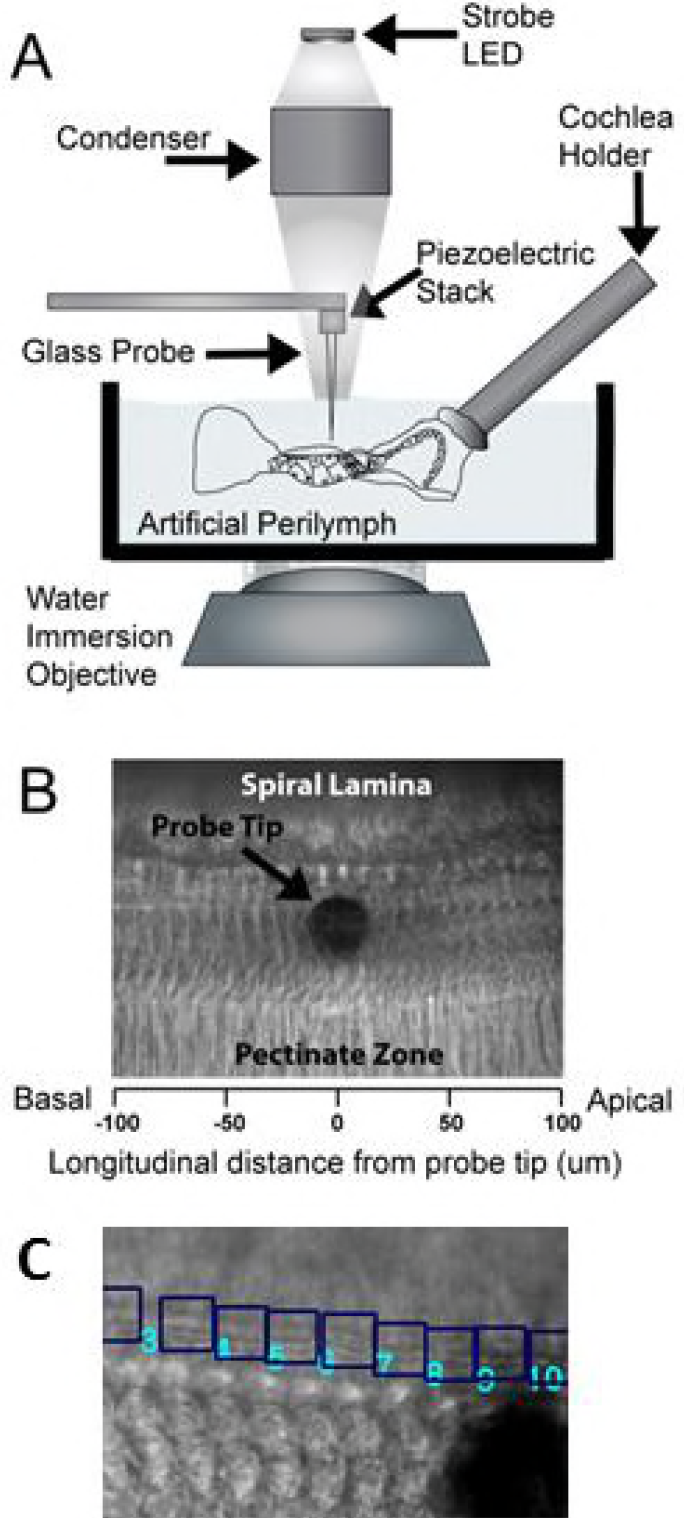
Panel A is an illustration of the experimental setup. The cochlea preparation is mounted to the stage of an inverted microscope. It is submerged in artificial perilymph in a dish with a cover slip bottom to maximize image fidelity. The probe assembly was made as small as practical since it is in the condenser light path. Panel B is typical of images acquired during an experiment. The dark circle in the middle of the image is the shadow of the probe tip. It is normal to the basilar membrane plane. The radial collagen fibers can be seen in the pectinate zone. Rows of outer hair cells are visible adjacent to the probe. Panel C is a magnified view of the IHCsc with their 64 × 64 pixel regions of interest (ROI) demarcated by the blue squares. The ROIs are 9.46 μm squares. The black circular shadow is the probe tip.

The stroboscopically illuminated microscope imaging system used was conceptually similar to the one developed by Karavitaki et al (27) for measuring electrically evoked outer hair cell displacement. In this study the transmitted light illuminator was retrofitted with a SML-LX1610USBC 350mW white LED (Lumex, Carol Stream, Illinois). A 20×, 0.5 NA or 40×, 0.8 NA water immersion objective (Olympus Waltham, Massachusetts) was used during all experiments. An additional 1.5× tube lens was used with the 20× objective to increase the total magnification tp 30×. The field of view of the 20× objective data was 315 μm in the longitudinal dimension, and 236 μm in the radial dimension. For the 40× objective it was 236 μm longitudinal and 177 μm radial. The entire radial width of the basilar membrane (160 μm) could be imaged in the turn 1 preparations. In the turn 2 preparations the window in the bone could not be made large enough to view the spiral ligament without causing damage. A PL959 (Pixelink Rochester, NY) black and white 1600 × 1200 pixel CCD USB camera was used to capture images. The white LED illuminator was driven by a custom made computer-controlled current amplifier. Low magnification images were used to identify probe distance from the base of the cochlea.

The tissue was stimulated with a pulled glass micropipette blunted to 25 - 45 μm in diameter with a microforge. The pipette was bonded with West System 105 epoxy thickened with colloidal silica (Gougeon Brothers, Bay City, Missouri) to an AE0203D04F piezoelectric stack actuator (Thorlabs, Newton, New Jersey). The probe assembly was designed to minimize its size to prevent optical interference due to its position within the optical path of the light condenser. The probe assembly was mounted to the motorized micromanipulator attached to the stage of the inverted microscope. During experiments the probe was mounted with the pipette normal to the microscope stage and thus the focal plane. The piezo stack was driven by a modified audio amplifier (Norelco-Philips, Andover, Massachusetts) connected to the output of a purpose built fourth order low pass anti-image filter that was connected to a PCI-6110e (National Instruments, Austin, Texas) digital to analog converter. The magnitude change and phase delay attributed to the low pass filter was processed out of the data using calibrations of probe displacement. The microscope was enclosed in a light blocking enclosure during experiments to attenuate noise from outside light sources. During experiments the peak to peak transverse displacement was chosen to be between 400 and 800 nm. In this range there was sufficient signal to noise ratio while maintaining the CP displacement in a physiologically relevant range. The displacement of the probe was calibrated using OHV 511 laser Doppler velocimeter (LDV) (Polytec, Hudson, Massachusetts).

The camera, strobe illuminator, and the stack actuator were all driven and controlled by a computer with a National Instruments data acquisition card. Custom software written in C# (Microsoft Corp., Redmond, Washington) and MATLAB (Mathworks, Natick, Massachusetts) automated the experiments and organized the image and stimulus data. During stimulus presentation, the data collection program opened the electronic CCD camera shutter, presented a sinusoidal stimulus waveform to the probe amplifier, strobed the illuminator with a 10% duty cycle pulse train that was phase locked to the stimulus waveform, then closed the shutter. The acquired image was saved and the process was repeated with the strobe signal advanced 45 degrees in phase. Eight images each representing a different phase were captured for frequencies ranging from 10 Hz to 42.5 kHz, producing an eight frame sequence at each frequency. Data was collected at multiple focal planes. In this study the inner hair cell stereocilia data was considered.

### Image processing

In plane displacement of the stereocilia was measured optically using a cross-correlation algorithm derived from Karavitaki (27) that computed displacement of feature edges in successive image phases in a selected region of interest (ROI). The stereocilia of each inner hair cell were tagged by hand within a 64 × 64 pixel ROI, An 11 × 11 pixel Gaussian low pass filter with a sigma of 1 was applied to each image. The images were Sorbel filtered to enhance the edges and remove low spatial frequency information. Each ROI was then extracted out of the full image and windowed with a raised cosine filter with a taper constant of 0.07 to attenuate spectral leakage. The resulting ROI was transformed into the frequency domain with a two dimensional Fast Fourier Transform. The original ROI size of 64 × 64 pixels was then increased to 1024 × 1024 by padding with zeros around the perimeter. The zero padding in the Fourier domain increased the spatial resolution of the images, enabling sub pixel resolution. Each phase’s padded ROI images were cross-correlated in the Fourier domain with each of the following phase images of the same ROI to find the displacement in pixels. The location of the maximum value in the cross-correlation matrix corresponded to the displacement of the features in the ROI. The displacement time progression of all 8 phases including cross correlating the zero phase image with itself reconstructed one cycle of a particular sinusoidal stimulus frequency displacement. The displacement time progression was fast Fourier transformed to extract magnitude and phase of the displacement for each ROI. The image pixel size was calibrated using a 2280-15 micrometer slide (Ted Pella Inc. Redding, California). The algorithm was calibrated by measuring the displacement of the calibrated probe.

## RESULTS

A preparation was considered successful and included in the analysis if there was no visible damage to the CP, spiral ligament, and inner osseous lamina under microscope observation within 1000 μm of the probe location. In all results shown the displacement of the stereocilia was normalized to the magnitude and phase of the probe displacement.

### Observations of mechanically-induced forward and reverse traveling waves in the cochlea

Figure 3 shows data from two typical experiments: One at the turn 1 longitudinal location (basal), and one at the turn 2 (more apical than turn 1) location. The turn 1 location has been estimated to be most sensitive (CF) at 34.2 KHz and the turn 2 is estimated to be most sensitive at 2.5 kHz (29). The radial displacement magnitude and phase of IHCsc at several frequencies are plotted as a function of longitudinal distance from the probe tip at both turn 1 and turn 2 longitudinal measurement locations.

**Figure 3.**
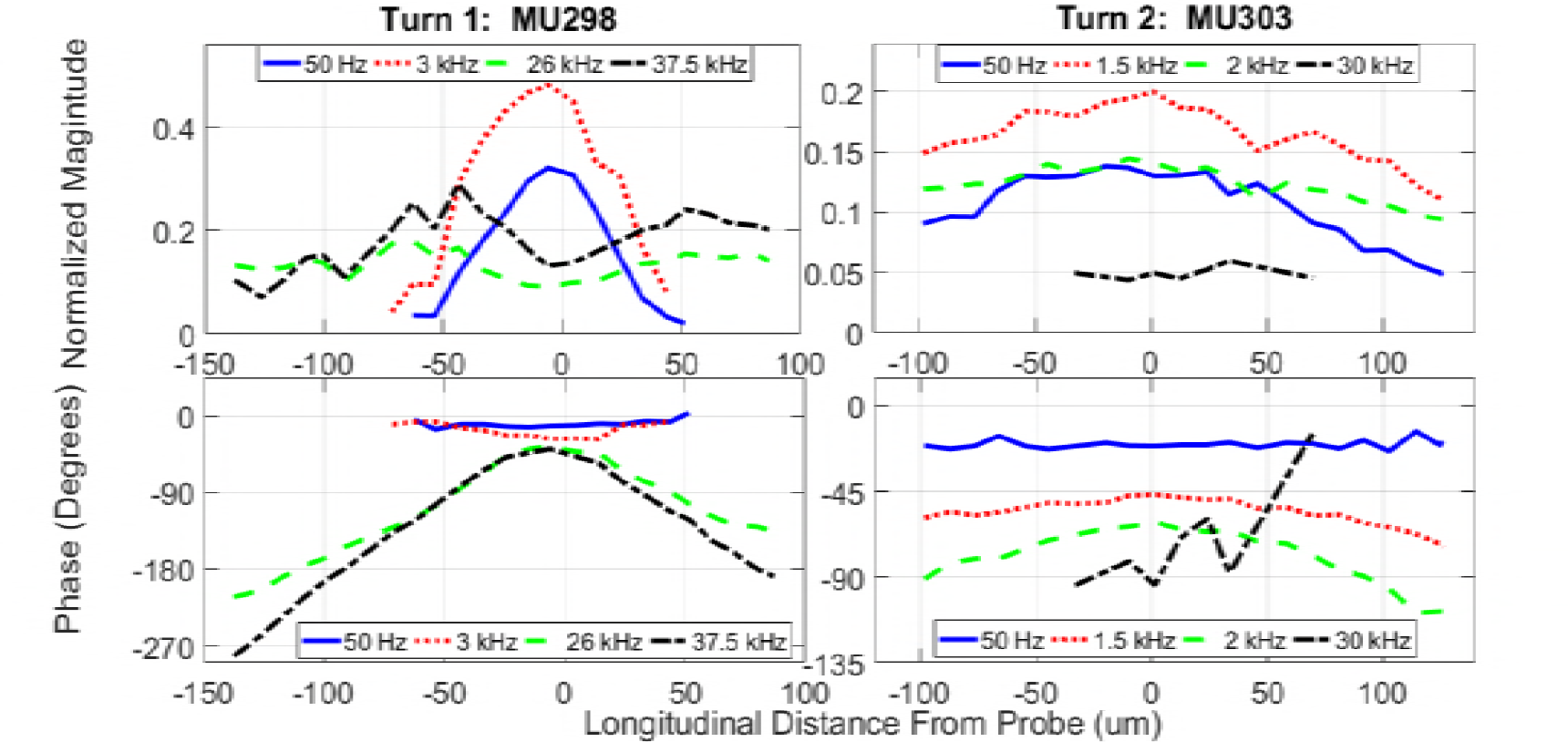
Radial displacement of IHC stereocilia basal and apical to the probe tip. Apical longitudinal distance from the probe is positive, basal is negative. Left panels are magnitude and phase at four frequencies from a turn 1 preparation. The probe is 1325 μm from the basal end of the cochlea corresponding to an estimated CF of 34.2 kHz (29). The right panes are magnitude and phase at four frequencies in a turn 2 preparation. The probe is estimated to be 6815 μm from the basal end of the cochlea corresponding to an estimated CF of 2.5 kHz (29). For both experiments the probe transverse displacement magnitude was 800nm. Points less than 8 nm in magnitude have been omitted because they are below the noise floor.

In turn 1 preparations, stimulated at frequencies well below CF (red and blue lines), the magnitude basal and apical to the probe shows a relatively narrow peak and decays into the noise floor (8 nm) within the field of view of the images. At these low frequencies there is minimal phase change over the measured longitudinal distance from the probe with respect to longitudinal location, indicating that the cochlear partition moves in phase over the observed distance. As the stimulus frequency increases (green and black lines), the peak of the magnitude broadens and a phase delay builds at the probe location. As the stimulus frequency approaches the CF, the magnitude persists for the entire field of view and the phase delay increases nearly linearly with distance from the probe. The decaying phase is indicative of traveling waves, propagating towards apical locations (in the forward direction) and basal locations (in the reverse direction) from the probe location.

In the second turn preparations similar observations are made compared to turn 1. At frequencies well below the CF (blue line) similar characteristics of the magnitude are observed, although the normalized peak is generally smaller. Although the optical field of view ends before the magnitude decays into the noise floor, we observe a peak at the probe location and a phase that is independent of the measurement locations. As the stimulus approaches the CF (red and green lines), the magnitude increases and the peak broadens. The phase delay at the probe increases as the frequency is increased. Similar to turn 1, traveling waves are observed for frequencies close to the CF, where the phase is accumulating at measurement locations further away from the probe. In turn 2 preparations the spatial phase delay slope at CF frequencies increases while moving both basal and apical from the probe. At frequencies higher than CF (black line) the magnitude decreases considerably and the phase becomes more random, suggesting the signal is in the noise floor.

### Turn 1 results

Figure 4 shows pooled data at multiple frequencies from 11 turn 1 experiments. The estimated CFs for these experiments range from 29 kHz to 40 kHz. The magnitude and phase are normalized to the probe displacement. At zero distance from the probe the data can be interpreted as a transfer function of the IHCs directly above the probe relative to the probe itself. Moving longitudinal to the probe the data can be interpreted as a spatial transfer function that shows the instantaneous phase.

**Figure 4.**
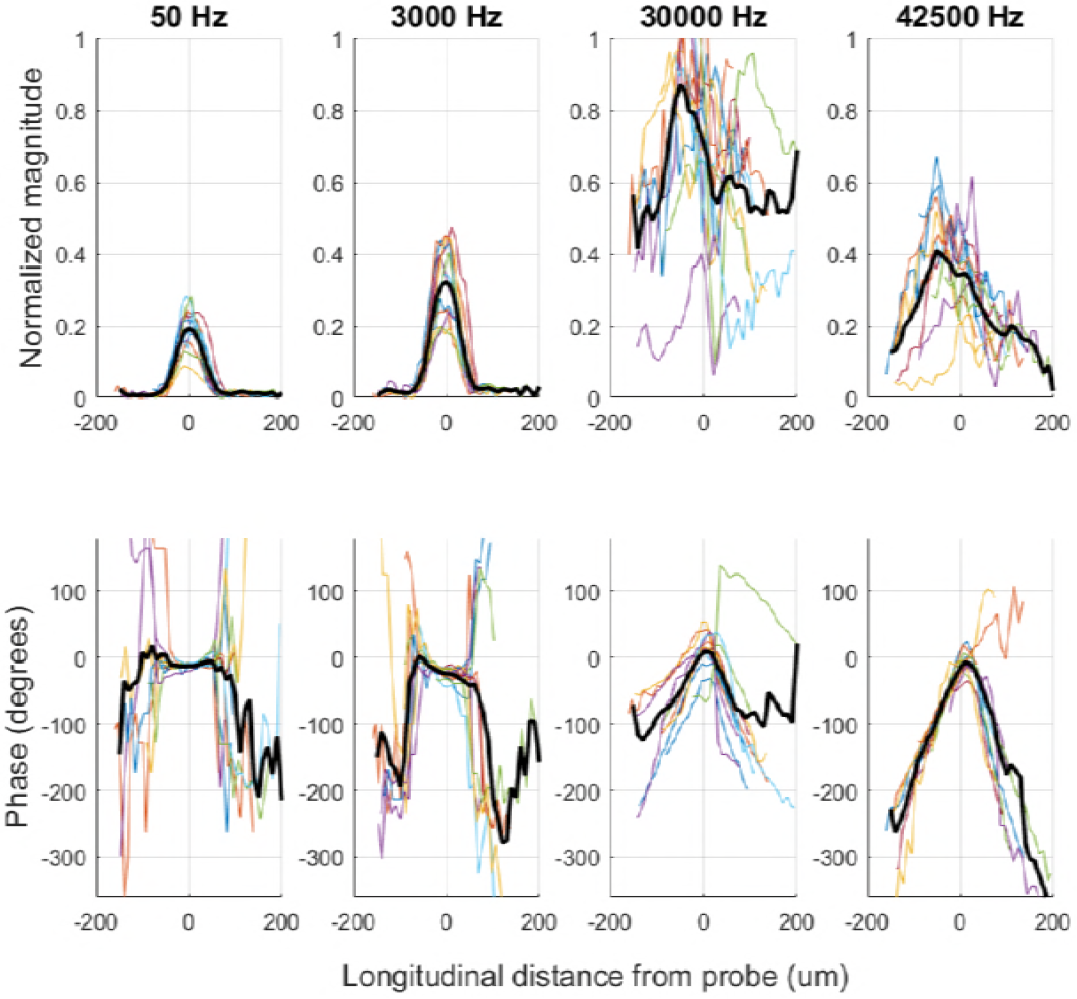
Magnitude and phase of pooled data from all turn 1 experiments. The four frequencies show the progression of the spatial response as the stimulus approaches estimated (29) CF (29 kHz to 40 kHz). The black traces are the average of the data from 11 animals. The lighter color traces are individual experiment responses. Frequency Responses are normalized to the calibrated probe frequency response.

At frequencies much below CF (50 Hz) a narrow peak of the magnitude is observed, in the close vicinity to the probe location. At 50 Hz, the maximum value of the peak is 0.19 ± 0.055; the phase delay in the peak region is 12.8 ± 3.1 degrees. At locations basal and apical to the probe, the magnitude decays into the noise floor within ±50 μm of the probe location. The phase remains constant within ±50 μm of the probe before it becomes random due to the noise floor. As the stimulus frequency is increased to 3000 Hz, the magnitude peak at the probe location increases to 0.32 ± 0.1 and the phase delay increases to 24.1 ± 8.9 degrees in the vicinity of the probe. Overall, the results for a stimulus freuqency of 3000 Hz appear very similar compared to the results reported for 50 Hz. Near CF (30000 Hz), the peak value of the magnitude increases significantly to 0.7 ± 0.19. The phase at the probe location is 7.9 ± 30.7 degrees, indicating more variability comared to the below-CF frequencies. Near CF, there is a consistent trend across experiments of an increase in phase delay both apical and basal to the probe. The phase accumulation at more apical locations is indicative of forward traveling waves and the phase delay at basal locations suggests reverse traveling waves. Lastly, as the stimulus frequency increases past CF (42500 Hz), the magnitude decreases significantly with a peak of 0.34 ± 0.1 basal to the probe. Although no significant phase delay is observed at the probe tip, the phase accumulations at apical and basal measurement locations suggest forward and reverse traveling waves for this frequency.

### Turn 2 results

Figure 5 shows the results of turn 2 data pooled from 7 turn 2 experiments. For all the turn 2 data the CFs are estimated to be approximately 2 kHz to 3 kHz.

**Figure 5.**
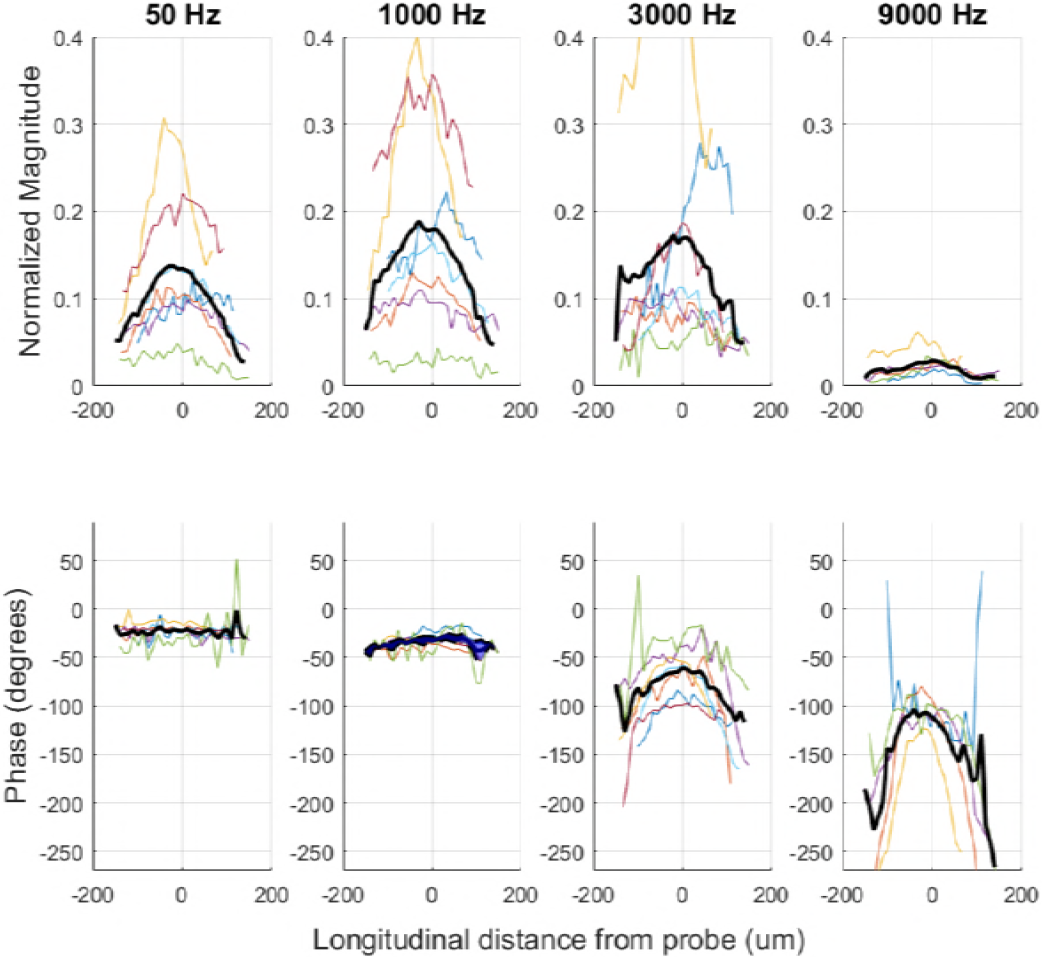
Magnitude and phase of pooled data from all turn 2 experiments. The four frequencies show the progression of the spatial response as the stimulus frequency approaches the estimated (29) CF (2kHz to 3kHz). The black traces are the average of the data from 7 animals. The lighter color traces are individual experiment responses. Frequency Responses are normalized to the calibrated probe frequency response.

At frequencies well below the CF (50 Hz) the magnitude has a broader peak than turn 1 data but a similarly flat phase. At the probe location the magnitude peak is 0.15 ± 0.09 and the phase amounts to −22 ± 6.7 degrees. Whereas the magnitude decays at more basal and apical measurement locations with respect to the probe, the phase stays constant. The signal remains above the noise floor in the image field of view therefore the phase remains coherent. At a 1 kHz stimulus frequency the average magnitude increases to 0.18 ± 0.12 at the probe, although not significantly. The phase delay increases significantly to −32 ± 8 degrees and a spatial delay starts to develop moving away from the probe. As the frequency increases to 3 kHz the average magnitude at the probe is 0.17 ± 0.15 and the phase delay is −61 ± 25 degrees at the probe. The spatial phase delay moving away from the probe increases. At 9 kHz the peak of the magnitude is 0.03 ±0.01 and the phase delay is −112 ± 15.5 degrees at the probe tip. The magnitude is significantly diminished above CF and the phase delay at the probe increases as is the spatial delay moving away from the probe.

Significant differences between turn 1 and turn 2 data can be seen. In turn 2 experiments, the magnitude moving away from the probe decreases slower compared to turn 1 experiments, indicating a longer longitudinal coupling. The phase delay at the probe location in turn 2 experiments increases with frequency whereas it does not in turn 1 experiments. Near CF both locations show an increasing spatial phase delay moving away from the probe. In turn 1 experiments this delay is more linear within the field of view than in turn 2 data.

### Evidence of forward and reverse traveling waves near the characteristic frequency

In both turn 1 and turn 2 preparations, near the respective CFs there is a spatial phase delay that builds moving further from the probe location. This suggests presence of both a forward (apical) and reverse (basal) traveling wave propagating from the source (the probe stimulus).

Traveling wave velocity and wavelength were computed at both longitudinal locations at frequencies near the CF, where there was evidence of a traveling wave. The slope of the phase vs. position was fit to a straight line by using the robust fit function in MATLAB (30). Wavelength (λ) was computed as: λ=2π Δx/Δphase. Traveling wave velocity was computed by multiplying the wavelength by stimulus frequency. Wavelength at frequencies much below CF cannot be calculated because they are either nonexistent in this preparation, or the wavelengths are too long to measure any appreciable phase delay within our field of view.

### Spatial decay of magnitude at low frequencies

For both turns the decay rates of the magnitude at low frequencies were fit to a Gaussian function (1) using MATLAB. Where *c* is defined as the space constant and is the distance when the magnitude is 1/e times or 37% of the peak value (where the probe tip was located). This measure has been used by others to quantify longitudinal coupling (31–33). Space constants were calculated at frequencies lower than one decade below the estimated CF (figure 6) where there was no significant accumulation of phase delay. At 10 Hz the space constant was 37.7 μm ± 5.44 μm in the base (N=13) and 141 μm ± 29.4 μm in second turn (N=7).

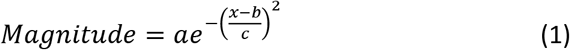

**Figure 6.**
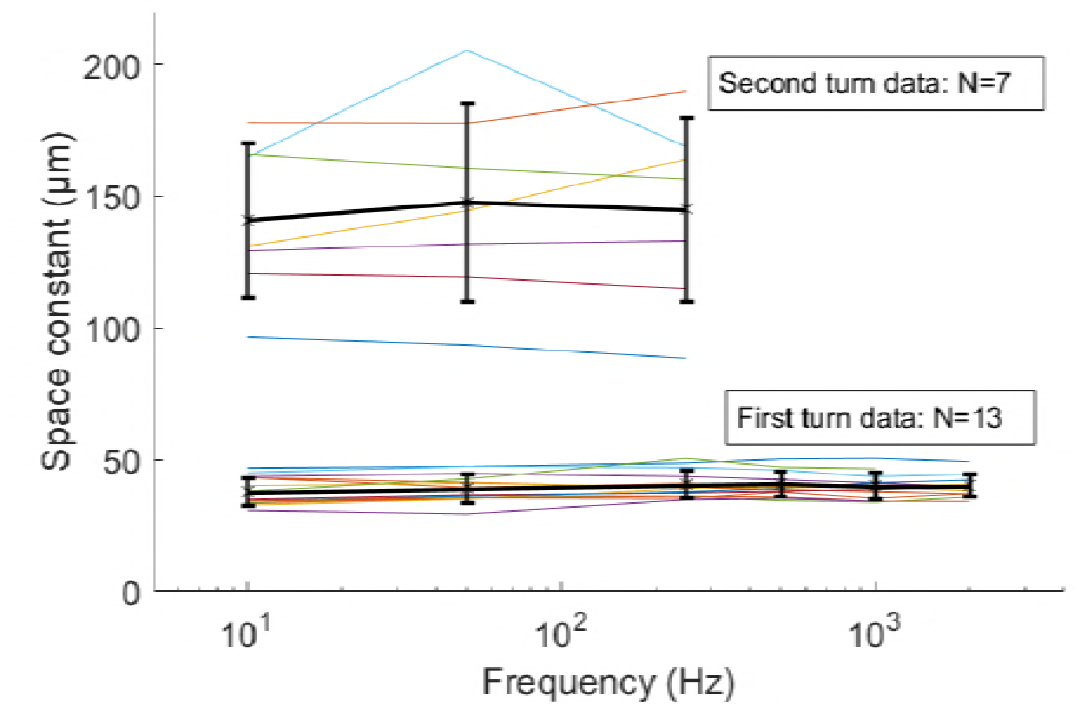
Plot of computed space constant ‘c’ vs. frequency for turn 1 and 2 preparations. Thin lines are individual experiment results, thick lines are the average with standard deviations.

The wavelength fraction, defined as the space constant divided by the measured traveling wavelength, was computed for both longitudinal locations. The results are plotted in table 1. For both locations the result was 14 - 22% indicating that low frequency coupling was significantly less than a traveling wavelength near CF.

**Table 1.** Measured traveling wave parameters. The wavelengths and velocities were calculated from phase slope data at stimulus frequencies near the estimated CF of the longitudinal place. For the turn 1 data, the basal and apical slopes from 27.5 to 40 kHz were averaged and standard deviation was calculated. The same procedure was performed on turn 2 data for stimulus frequencies from 2 to 3 kHz. The space constant was calculated from the spatial decay of the 10 Hz data for each turn. Space constant data is a composite of basal and apical data. The wavelength fraction is the space constant divided by the wavelength.

|  |  | Wavelength<br>1/k | STD | TW Velocity | STD | Space constant | SC / k<br>Wavelength fraction |
| --- | --- | --- | --- | --- | --- | --- | --- |
| Turn 1<br>CF 29 kHz –<br>40 kHz | Basal | 244 $\mu\text{m}$ | 58 | 8.0 m/s | 0.7 m/s | 37.7 $\mu\text{m}$ | 0.15 |
| | Apical | 174 | 23 | 5.9 | 1.4 | 37.7 $\mu\text{m}$ | 0.22 |
| Turn 2<br>CF 2 kHz -3<br>kHz | Basal | 978 | 239 | 2.4 | 0.2 | 141 $\mu\text{m}$ | 0.14 |
| | Apical | 787 | 248 | 1.9 | 0.2 | 141 $\mu\text{m}$ | 0.18 |

## Discussion

### Presence of traveling waves in the cochlear partition

In both measurement locations at frequencies where traveling waves were present, there were both forward and reverse waves in the CP. The variability in the data at CF and greater frequencies may be in part due to the difference in CF of the individual experiments (29 kHz to 40 kHz for turn 1 and 2kHz to 3kHz for turn 2). The presence of the forward traveling wave mode in the CP was expected from the large body of work that predicted and measured them (5–7, 9, 18, 34, 35). The measurement of the expected forward traveling wave in our results improves the confidence that the measured reverse traveling wave is not an artifact. Our measured traveling wave length result near CF measured at the turn 1 location (table 1) is comparable to Ren (35), although they report only a forward traveling wave. The experiments differ in how the BM is stimulated. In Ren the preparation was alive and driven from the stapes. In our preparation the basilar membrane was driven by a point source directly on the BM. Our preparations were excised cochleae where we can assume the active processes were severely diminished, if not absent. However, it should be noted that Karavitaki (27) demonstrated outer hair cell motility from electrical stimulation using a similar preparation and the same culture medium recipe. Measurements made by Ghaffari (36) in mouse tectorial membrane reported traveling wave velocities that were within a factor of two of our turn 1 data. Their data has the same trend of higher measured velocities in basal traveling waves than in apical traveling waves and lower velocities at lower frequencies. Temchin (9) reports similar velocities in chinchilla using measures of auditory nerve recordings. Like our results, they report a lower velocity at more apical (lower CF) locations.

De Boer (7), using a 3 dimensional model based on guinea pig data, predicts the presence of forward and reverse traveling waves that are created simultaneously in time at the generation site. Their model is replicated in our experiments where they stimulate a location along the basilar membrane with a 59 μm wide pressure source. This is similar in size to our probe tip. As seen in figure 2 of their paper (7), their model predicts forward and reverse propagating waves in the cochlear partition when the stimulus frequency is higher and more basal to the CF location. Their forward and reverse wavelength is approximately 4332 μm corresponding to a velocity of 73 m/s at 17 kHz. When their stimulus location and frequency are slightly apical to CF, the wavelength and velocity decrease to 1080 μm and 18 m/s respectively. Their result is consistent with this study where there are both forward and reverse traveling waves created simultaneously in time, and the wavelength and velocity decrease when the stimulus frequency is near the CF. Their model wavelengths and velocities are larger; that may be a result of the difference in species, or the model simplifications they describe in their paper. Our results could also be lower due to the open scalae.

It is unclear how far basal the CP reverse traveling wave mode persists. In both turn 1 and turn 2 experiments the traveling wave magnitude persisted within the field of view of the images. In the turn 1 data more than one wavelength was observed within the field of view. The field of view limits the ability to determine spatially where or if the reverse traveling wave magnitude decays into the noise floor. If reverse CP traveling waves contribute to otoacoustic emissions by driving the stapes they must result in a fluid pressure in the scala vestibuli suitable to drive the stapes footplate. Li (12) derived reciprocity in their cochlear model that suggests a CP velocity generated from an internal source, that is similar to our probe, can drive the stapes.

Active processes and structural impedance irregularities are both proposed to be sources of otoacoustic emissions (3, 4). If the probe is considered to be a reflection point at an impedance irregularity or displacement generated from an active process, our results provide insight into otoacoustic emission propagation modes. If impedance discontinuities are present in the CP, our results support the coherent reflection theory of Zweig and Shera (37) where time and space phase coherent discontinuities cause reflections that lead to reverse traveling waves. The width of our probe tip coupled to the basilar membrane provides a coherent driving pressure for the BM. The reflections in a healthy intact cochlea will result in smaller magnitude reverse traveling waves than in forward traveling waves due to the small difference in impedance that causes the reflection. If the mechanics of the cochlear partition are approximated as linear at low stimulus levels in experiments or models studying traveling waves, the large amplitude forward-propagating wave possibly obscures the reverse-propagating modes, making them difficult to measure in experiments. Bowling and Meaud (15) have demonstrated forward and reverse traveling waves as well as a compression wave are present in a model that features realistic active processes. They report that the distortion product otoacoustic emission-generated waves are dominated by the slow reverse traveling wave mode as opposed to the compression wave.

### Phase at locations near the probe

In the second turn experiments, as the stimulus frequency increases the phase delay of the RL relative to the BM increases at the RL location directly above the probe. At 50 Hz there is on average 21 degrees of delay. At predicted CF (2kHz-3kHz) there is on average 61 degrees of delay. The 40 degree difference is consistent with that seen by Chen. This result suggests that there is differential displacement of the RL and BM in the second turn. In their preparation they reported the RL leads the BM where we observe the BM leads the RL. The discrepancy may be due to the different methods of stimulation. Chen stimulated the oval window thus drove the CP with fluid pressure where we drive it in reverse by direct stimulation of the CP from the scala tympani.

In turn 1 the phase delay doubles from 12 degrees at 50Hz to 24 degrees at 3kHz. As the frequency approaches CF there is no significant phase delay. It is possible that the probe is buckling at higher frequencies and the unloaded probe calibration does not take this into effect. Based on measured mechanical stiffness by Naidu (38), the probe is likely presented with a higher impedance in turn one than in turn 2. The thinner pectinate zone (PZ) and shorter pillar cells may be stiffer, thus resisting deflection from the impedance loads of the scalae fluid, RL, and TM. BM stiffness measurements by Olson (39) and Naidu (38) suggest that the OoC contributes to the point stiffness of the BM in gerbil. Olson showed that the stiffness of the BM changes radially depending on what cellular structures are above the measurement location. Naidu showed that removing the organ of Corti reduced the point stiffness of the BM. Thus the OoC is expected to contribute to the impedance seen by the probe.

The phase at locations near the probe may also be affected by the arched shape of the pectinate zone in gerbil (40, 41). It is possible the probe tip is buckling the pectinate arch (figure 7), leading to the phase delay in the turn 2 data, although there is no conclusive evidence. The placement of the probe under the pillar cell minimizes the force on the arch by pushing against its attachment where the cross section is minimal. If arch buckling is occurring, it is possible that some of the variability across experiments is a result of static force that changes the geometry of the arch thus resulting in transfer function differences. Based on space constant measurements of the BM from Naidu (32) and Emadi (33), if the probe disturbed the mechanics of the BM, the effect will rapidly be attenuated moving longitudinal to the probe. Under this assumption the traveling wave velocity measurements of the IHCsc should not be affected.

**Figure 7:** Cartoon depiction of probe tip (P) indenting the arched basilar membrane. The static displacement caused by placing the probe on the basilar membrane may cause distortion in the cochlear partition that could affect the mechanics of the cochlear partition. Notably the collagen fibers in the Arcuate (AZ) and pectinate (PZ) zones may be disrupted. The outer pillar cell (OPC) may be pushed up by the probe causing the geometry to change.

### Longitudinal mechanical coupling

The presence of stereocilia displacement at low frequencies both apical and basal to the probe suggests that there is longitudinal coupling in the cochlear partition. The magnitude variablility at low frequencies may be due to differences in the probe contact with the cochelar partition. Under the assumption that at very low stimulus frequencies the effect of the scalae fluid and cochlear partition mass are negligible, then the stiffness and geometry of the structures that comprise the cochlear partition govern their displacement when subject to a displacement from the probe.

Experiments by Voldrich (42), where the BM was indented by a pin and the deflection pattern observed, indicated that there was no significant longitudinal coupling in the basilar membrane thus the basilar membrane can be treated as an array of independent resonators. Naidu (32) demonstrated that there was significant longitudinal coupling in the BM. Naidu reported a coupling space constant of 16.7 μm ± 1.9 μm in the basal turn and 34.5 μm ± 8 μm in the second turn. Emadi (33) reported a space constant in second turn of gerbil of 20.6 μm ±7.6 μm. As stated by Emadi (33), a space constant of approximately 20um is on the order of two hair cell diameters indicating little longitudinal coupling. Newburg (43) measured the space constant of lateral displacement of coupling in the PZ radial fibers to be 3.59 μm.

Our results indicate a significantly longer space constant than Voldrich (42), Naidu (32), Emadi (33), and Newburg (43). The most likely explanation for the difference is that our data is observing coupling in the reticular lamina by measuring hair bundle displacement, not in the pectinate zone collagen fibers. The increase in space constant is consistent with the RL acting as a plate that is in shear and bending as a result of transverse displacement of the BM being coupled to the RL via the outer pillar cell head. The shear could be a result of adjacent pillar cells and their connection to the basilar membrane providing a restoring force. Although our space constant measurements are longer than other measurements, they are shorter than the wavelengths of traveling waves measured at CF for both locations. Our calculation of the wavelength fraction, where the space constant is divided by the traveling wave length, indicates that longitudinal coupling is less than 1/4 of a wavelength at CF.

### Effects of open scalae

During the experiments both scalae were open which would expectedly result in attenuation of the fast fluid pressure compression wave that is propagates nearly instantaneously (approximately 1500 m/s) down the length of the basilar membrane in an intact cochlea. Based on models from Steele and Zais (1985), even with open scalae there will be a slow pressure wave at the top and bottom boundary of the CP, if the fluid depths in the scalae are great enough to mass load the motion of the cochlear partition. Their results suggest that maintaining greater than 1/10 of the original fluid depth results in a minimal effect on frequency peak location. The preparations used in these experiments were immersed in culture medium that resulted in a much deeper fluid depth than in an intact cochlea. The fluid depth was greater than 1mm in both the ST and SV sides of the preparation. This is greater than the traveling wavelengths measured in both turn 1 and turn 2 experiments.

Results from Patuzzi (44), where they opened and drained the scala tympani showed large losses of sensitivity but indicated no change to the CF suggesting that measurements made with open scalae are valid for making passive measurements. The fact that the sensitivity decreases but the frequency response has a similar characteristic suggests that the hydromechanical elements that comprise the CP are maintained. The draining of the cochlea results in a reduction in mass loading of the CP. Experiments by Olson (45), where intracochlear pressure measurements were made through small holes drilled in the scalae, indicate that the pressure peaks generated by the CP-fluid interaction are attenuated as the pressure sensor moves away from the CP. This supports the result of Steele and Zais (46), where fluid depth is more critical than a closed scalae to maintain a physiologically relevant preparation. Results by Ulfendahl (47) in guinea pig cochleae demonstrated that the mechanical response of the basilar membrane was attenuated at low frequencies (300Hz) when SV was opened.

Their results suggest that the mass loading of the scalae fluid contributes more to higher frequency tuning than a sealed cochlea. It is likely in our experiments that the open scalae disturbed the tuning but the characteristic behavior of the system was maintained. The lack of measureable traveling waves at frequencies lower than CF is likely due to the open scalae and the wavelengths being too long to resolve within our field of view.

## Conclusion

Our data suggests that forward as well as reverse traveling wave modes do exist in the structure of the CP. On this basis it is possible that local impedance mismatches in the CP will result in reverse propagating traveling waves by means of reflections. In addition, active processes from the OHCs may reverse propagate along the CP. These reverse propagating waves may contribute to otoacoustic emissions.

The longitudinal coupling space constants suggest that IHCs adjacent to a CF location are also activated when a CF location responds to a tone. Similarly, the OHCs adjacent to a CF location will be displaced and may contribute to force generation. The spatial recruitment of more OHCs at a location may result in a more favorable energy transfer from the OHCs to couple their energy into the fluid in the scalae.

The computed wavelength fraction suggests that the longitudinal coupling in the CP has a significant effect on the wavelengths of the traveling waves that a particular location along the cochlea can support. Therefore, both point impedance and longitudinal coupling mechanics of the CP at a particular location are interdependent.

## Author Contributions

AZ and LCR performed the research and data analysis. DCM was the principal investigator. All authors contributed to the manuscript.

## Acknowledgements

David J. Anderson, Andrew Brughera, Allyn E. Hubbard, Heidi H. Nakajima, Stefan Raufer.

This research was funded by the National Institutes of Health (NIH) # R01 DC000029.

